# Creating DNAm Algorithms Using the Illumina Methylation Screening Array (MSA)

**DOI:** 10.64898/2026.08.31.748429

**Authors:** Kirsten B Seale, Sayf Hassouneh, Ilinca Giosan, Karen Sugden, Laura Balagué-Dobón, Varun B. Dwaraka, Jessica Lasky-Su, Mike Mallin, Avshalom Caspi, Terrie Moffitt, Ryan Smith, Natàlia Carreras-Gallo

## Abstract

Most established DNA methylation (DNAm) biomarkers were developed on legacy Illumina EPIC arrays. The Infinium Methylation Screening Array (MSA) offers a lower-cost, higher-throughput alternative with reduced probe content, but EPIC-trained algorithms cannot be assumed to transfer directly. Here we present a reproducibility-based framework for developing and transferring DNAm algorithms on the MSA. Using paired biological replicates profiled on EPICv1 and MSA (1,764 EPICv1–MSA sample pairs, plus within-array MSA replicates on the same and different beadchips), we quantified probe-level agreement using mean absolute error (MAE) and intraclass correlation coefficients (ICC). Of 140,150 CpG sites shared between EPICv1 and MSA, 40,786 (29.1%) met both stability criteria (MAE < 0.05 and ICC(2,k) > 0.6). This stable feature space supported two modelling streams. First, we trained 134 epigenetic biomarker proxies (EBPs) natively on MSA, with and without kernel principal component analysis (kPCA) for sample-level harmonisation. All 134 reached same-beadchip ICC(2,1) ≥ 0.80 (median 0.97) and 96.3% reached different-beadchip ICC(2,1) ≥ 0.60 (median 0.81), with a median Spearman correlation of 0.48 against observed values. Among the 72 kPCA-selected models with a comparable stable-probe baseline, 70 (97%) showed higher cross-beadchip ICC (median improvement +0.18). Second, we transferred three established clocks using model-specific strategies: OMICmAge and SystemsAge were retrained to estimate their EPICv1-derived values (held-out test-set rho = 0.944 and 0.912–0.949), whereas DunedinPACE required stable-probe normalisation and robust linear calibration, which raised cross-array ICC(2,1) from 0.784–0.810 to 0.891–0.925 and reduced MAE from 0.085–0.089 to 0.041–0.050 across three sample sets. Reduced probe content does not preclude reproducible DNAm biomarker measurement, and transfer strategy must be matched to model architecture.

## Introduction

DNA methylation (DNAm), the covalent addition of a methyl group to cytosine residues at CpG dinucleotides, is a key epigenetic mechanism involved in the regulation of gene expression and maintenance of cellular identity. Over the past decade, DNAm-based biomarkers have emerged as powerful tools for quantifying aging and disease biology, expanding from early chronological age predictors to include measures of biological age, pace of aging, organ-system dysfunction, disease risk, and diverse circulating clinical and metabolomic traits ^1–5^. Their utility stems from the relative stability of DNAm patterns, their ability to integrate cumulative genetic and environmental influences, and the feasibility of generating measurements reproducibly at scale from minimally invasive blood samples ^6,7^. Consequently, DNAm algorithms are increasingly being incorporated into research and clinical workflows for risk stratification, longitudinal monitoring, and intervention assessment. However, most established DNAm clocks and biomarker panels were developed using legacy Illumina methylation arrays, creating challenges for their deployment on newer platforms.

Transitions between Illumina methylation arrays represent a major obstacle to the widespread implementation of DNAm biomarkers. Platforms such as the Infinium MethylationEPIC (EPICv1), EPICv2, and the recently introduced Methylation Screening Array (MSA) differ substantially in probe content, genomic coverage, and assay design ^8^. As a result, probe overlap is incomplete, and even shared CpG sites may not exhibit equivalent measurement characteristics across platforms. These differences can introduce technical variation that propagates into downstream models, reducing reproducibility and potentially altering biological interpretation ^9–13^. The challenge is particularly relevant for the MSA, which was designed as a lower-cost and more scalable platform for high-throughput and translational applications. While its streamlined design offers substantial advantages for large-scale deployment, many existing DNAm algorithms depend on CpGs that are absent or behave differently on the MSA platform, necessitating retraining, recalibration, or alternative adaptation strategies.

Despite growing interest in cross-platform deployment, existing approaches remain limited and fragmented. Prior studies have largely relied on probe-overlap filtering, direct application of legacy models, or global normalization procedures, often without systematically evaluating probe-level reproducibility or accounting for residual technical variation across platforms ^13,14^. Furthermore, most approaches implicitly assume that all DNAm models can be transferred using a common strategy, despite fundamental differences in model architecture and biological targets. Consequently, there is currently no unified framework for translating DNAm biomarkers across array platforms while preserving both technical validity and biological interpretability. Addressing this challenge requires approaches that jointly consider reproducible feature selection, mitigation of platform-specific technical effects, and the distinct transfer requirements of different classes of DNAm algorithms.

A central component of such a framework is the identification of CpG sites that are consistently and reliably measured across platforms. Shared CpGs cannot be assumed to be interchangeable, as technical discrepancies may distort both absolute methylation levels and relative biological signal. However, restricting analyses to reproducible probes alone may be insufficient, because residual platform- and beadchip-related effects can introduce nonlinear variation that degrades model performance. Moreover, different biomarker classes impose different constraints on transferability. Epigenetic biomarker proxies (EBPs) trained to predict clinical or molecular traits may be amenable to retraining using reproducible features, whereas train-to-estimate clocks and scaling-dependent algorithms may require distinct calibration or normalization procedures to preserve their measurement properties. Together, these considerations motivate a framework that integrates reproducibility-informed feature selection, sample-level harmonization, and model-specific transfer strategies.

In this study, we develop and validate a generalizable framework for transitioning DNAm algorithms from legacy EPIC-based platforms to the Illumina Methylation Screening Array. Using paired EPICv1 and MSA samples, we identify reproducible CpG features based on complementary measures of technical agreement and use this stable feature space to support multiple modelling approaches. We apply this framework to the development of 134 MSA-compatible EBPs, evaluate the impact of stable-probe feature selection and nonlinear harmonization on model performance, and implement tailored transfer strategies for established epigenetic clocks, including OMICmAge ^5^, SystemsAge ^15^, and DunedinPACE ^3^. Collectively, our results establish the MSA as a scalable platform for DNAm biomarker deployment and provide a practical roadmap for reproducible cross-array translation of epigenetic algorithms.

## Results

### Study design

This study was designed to transition existing DNAm epigenetic clocks from the legacy Illumina arrays (EPICv1) to the MSA array, while preserving cross-array reproducibility, as well as build novel MSA epigenetic biomarkers. Paired EPICv1 and MSA samples (Supplementary Table 1) were used to define a stable probe set for EPICv1 - MSA and MSA - MSA, based on a combination of mean absolute error (MAE) and intraclass correlation coefficient (ICC) thresholds (Table 1, Supplementary Table 2).

**Table 1.** Identification of Stable CpG Probes Across Arrays Using MAE and ICC Criteria.

| Array1 | Array2 | N Paired biological replicates | Num. CpGs shared | Num. CpGs MAE < 0.05 (%) | ICC(2,1) > 0.6 (%) | ICC(2,k) > 0.6 (%) |
| --- | --- | --- | --- | --- | --- | --- |
| EPICv1 | MSA | 1,764 | 140,150 | 103,595 (74%) | 32,474 (23%) | 55,437 (40%) |
| MSA replicates (different beadchip) |  | 32 | 267,677 | 231,490 (86%) | 75,742 (28%) | 109,969 (41%) |
| MSA replicates (same beadchip) |  | 24 | 267,082 | 263,034 (98%) | 130,220 (49%) | 163,992 (61%) |

Cells highlighted in green indicate fraction of probes meeting threshold is >60%; yellow indicates fraction of probes meeting threshold is between 30% and 60%, and red indicates fraction of probes meeting threshold is <30%.

The stable probes supported two analytical streams (Figure 1). For EBP development, feature selection was informed by same-beadchip MSA technical replicates, and models were trained either on the stable probes directly or on kPCA-derived embeddings. Final models were selected based on Spearman rho thresholds and replicate-based ICCs to ensure cross-beadchip reproducibility. For epigenetic clocks, OMICmAge and SystemsAge were retrained on MSA using the same MSA stable probe universe derived from same beadchip MSA technical replicates, whereas DunedinPACE required a separate calibration procedure. This framework provides a robust and generalisable approach for deploying DNAm algorithms on the MSA platform.

**Figure 1.**
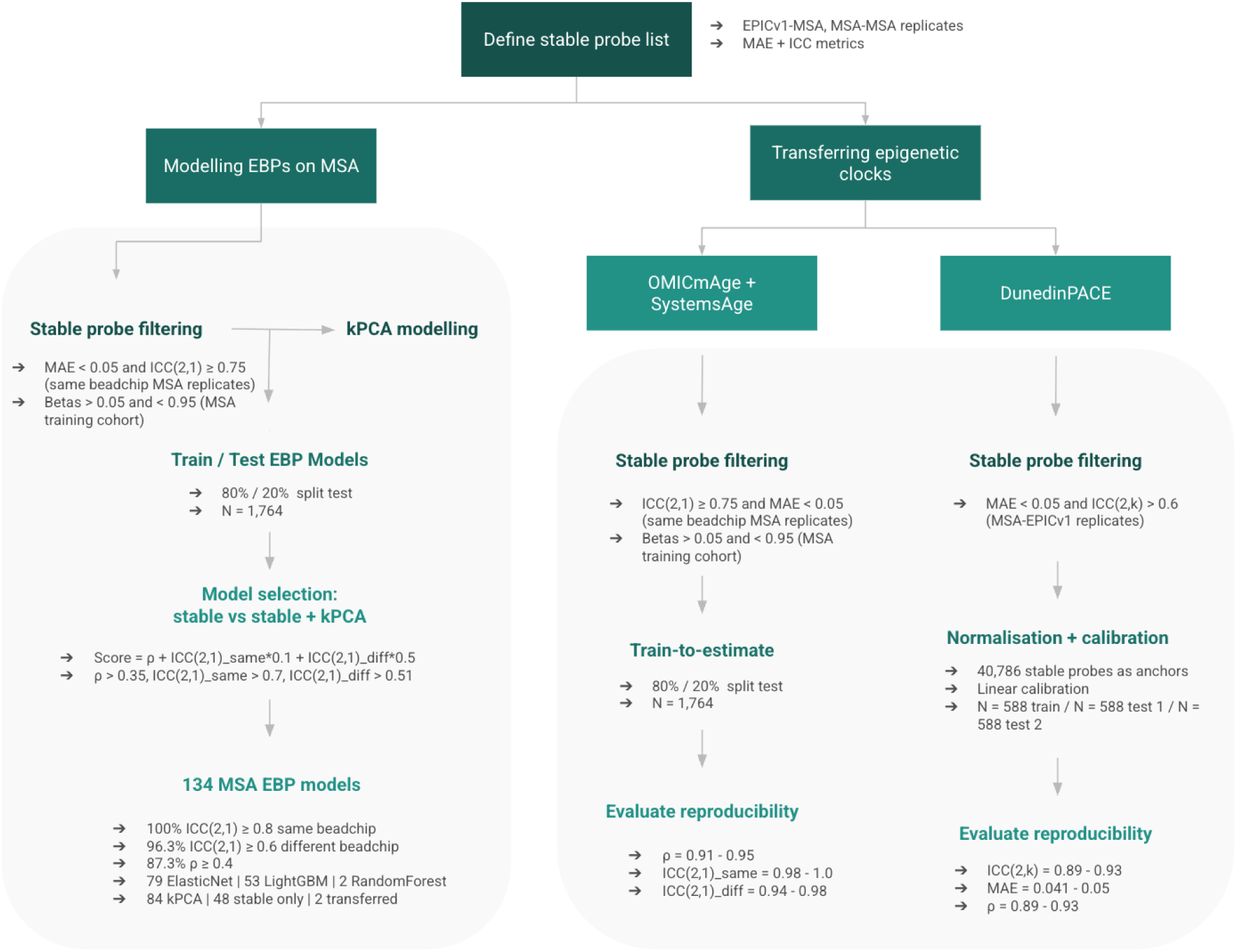
Study design for transitioning DNAm algorithms to the Illumina Methylation Screening Array (MSA). Schematic overview of the analytical framework. Paired biological replicates profiled on EPICv1-MSA and MSA-MSA arrays were used to identify reproducible CpG probes based on mean absolute error (MAE) and intraclass correlation coefficient (ICC) thresholds. The stable probe set supported two parallel modelling streams: (i) development of 134 epigenetic biomarker proxies (EBPs) using stable-probe and kernel principal component analysis (kPCA)-based approaches, and (ii) transfer of existing epigenetic clocks (OMICmAge, SystemsAge, and DunedinPACE) to the MSA platform.

### Identifying stable probes for cross-array feature selection

To define the high quality feature set for algorithm development, we used two complementary classes of probe-wise reproducibility metrics: MAE and ICC, computed on paired within-array and cross-array biological replicates (Table 1, Supplementary Table 1, Supplementary Table 2). MAE provides an absolute measure of disagreement by computing the average absolute difference in beta values between biological replicates. Because beta values have a natural 0–1 scale, this directly quantifies platform disagreement in methylation units and captures systematic offsets. In contrast, ICC quantifies the proportion of total variability attributable to true biological differences between samples rather than measurement noise.

We calculated both the ICC(2,1), a two-way random-effects, absolute-agreement model appropriate for generalizing across platforms, and the ICC(2,k), which evaluates the agreement of averaged measurements and reflects expected stability when multiple replicates contribute to an estimate (Supplementary Table 2). Using both MAE and ICC allowed us to jointly evaluate accuracy (low absolute differences) and consistency (high rank-order stability). Probes that scored well on both metrics (MAE < 0.05 and ICC(2,k) > 0.6) were classified as technically stable and exhibit consistent behaviour irrespective of array. ICC(2,k) defines the cross-array stable-probe set throughout, whereas ICC(2,1) is used to quantify single-measurement reproducibility of individual models and replicates.

Of the 140,150 CpG sites shared between EPICv1 and MSA with complete data, 40,786 (29.1%) met both stability thresholds, while 77,460 (55.3%) met one criterion (intermediate), and 21,904 (15.6%) met neither (unstable) (Figure 2A-C). The joint MAE-ICC classification system accurately stratifies probe quality, as illustrated by representative probes from each ICC class - Poor, Fair, Good, and Excellent - which show progressively tighter cross-platform agreement (Figure 2D-G). Cross-platform stable probes (EPICv1-MSA) were distributed evenly across genomic contexts, with a modestly higher stable fraction among Type II than Type I probes (30.7% vs 20.6%; Supplementary Figure 1). Probe-level MAE and ICC distributions are shown in Supplementary Figure 2.

**Figure 2.**
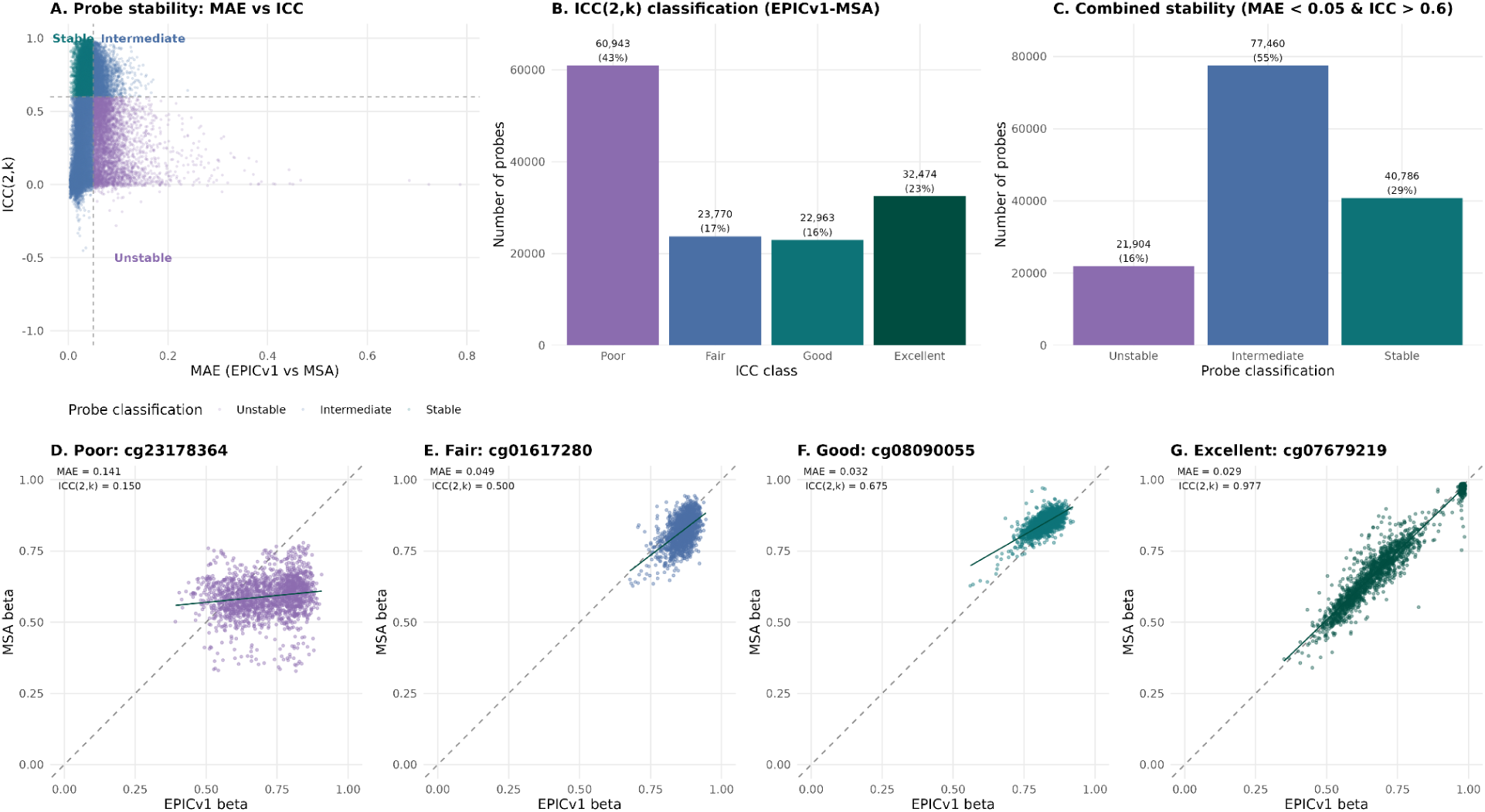
Cross-array probe stability between EPICv1 and MSA. **(A)** Scatter plot of MAE versus ICC(2,k) for 140,150 shared CpG sites with complete data. Probes are coloured by stability classification: stable (MAE < 0.05 and ICC(2,k) > 0.6; teal), intermediate (one criterion met; slate blue), and unstable (neither criterion met; purple). Dashed lines indicate classification thresholds. **(B)** Distribution of probes across four ICC(2,k) classes: Poor (< 0.4; n = 60,943; 43%), Fair (0.4-0.6; n = 23,770; 17%), Good (0.6-0.75; n = 22,963; 16%), and Excellent (>= 0.75; n = 32,474; 23%). **(C)** Distribution of probes across the three-tier stability classification: stable (n = 40,786; 29%), intermediate (n = 77,460; 55%), and unstable (n = 21,904; 16%). **(D-G)** Representative scatter plots of EPICv1 versus MSA beta values for individual probes from each ICC class, illustrating progressively tighter cross-platform agreement from Poor (D) to Excellent (G). Each point represents one paired sample (n = 1,764). Dashed line = identity; solid line = linear fit. MAE and ICC(2,k) values are annotated for each probe.

Stability was similarly assessed for the remaining array comparisons (Table 1). MSA within-array replicates were highly reproducible, particularly on the same beadchip (n = 24; 98% of shared probes with MAE < 0.05 and 61% with ICC(2,k) > 0.6), with a modest reduction across different beadchips (n = 32; 86% and 41%, respectively) that motivated the beadchip-aware harmonisation described below. Because MSA same-beadchip replicates additionally defined the feature space for EBP training, their high within-array reproducibility directly supports the modelling streams below.

These stability metrics underpin the two modelling streams: EBP development on MSA, and the transfer of OMICmAge, SystemsAge, and DunedinPACE to the MSA platform.

### Application of stable-probe and kPCA frameworks to MSA epigenetic biomarker proxies development

After establishing a set of technically robust CpG sites using reproducibility-based stable probe filtering, we evaluated how this feature set could be combined with nonlinear sample level harmonization to improve the performance of EBPs trained on the MSA platform. Restricting the feature space to stable probes substantially reduced measurement noise and yielded high cross-beadchip reproducibility for many biomarkers. However, several biomarkers continued to show diminished ICC(2,1) across beadchips, particularly those trained on smaller sample sizes or noisier endpoints, indicating persistent nonlinear array structure that could not be addressed through feature selection alone. To mitigate these effects, we incorporated kernel principal component analysis (kPCA) as an additional dimensionality reduction and harmonization step.

kPCA maps samples into a higher-dimensional feature space where platform-driven patterns can be more effectively separated from biological signals. Across models, kPCA consistently improved reproducibility in replicate comparisons, yielding same-chip ICCs exceeding 0.96 for most biomarkers and substantially improving cross-array ICC relative to both the original and stable-only models. Of the 84 kPCA-selected models, 72 had a comparable stable probe baseline; 70 of these (97%) showed improved different-beadchip ICC, with a median improvement of +0.18 (Figure 3). These improvements were especially pronounced for biomarkers with previously low reproducibility, confirming that kPCA provides a robust strategy for mitigating nonlinear batch effects that remain after probe-level filtering.

**Figure 3.**
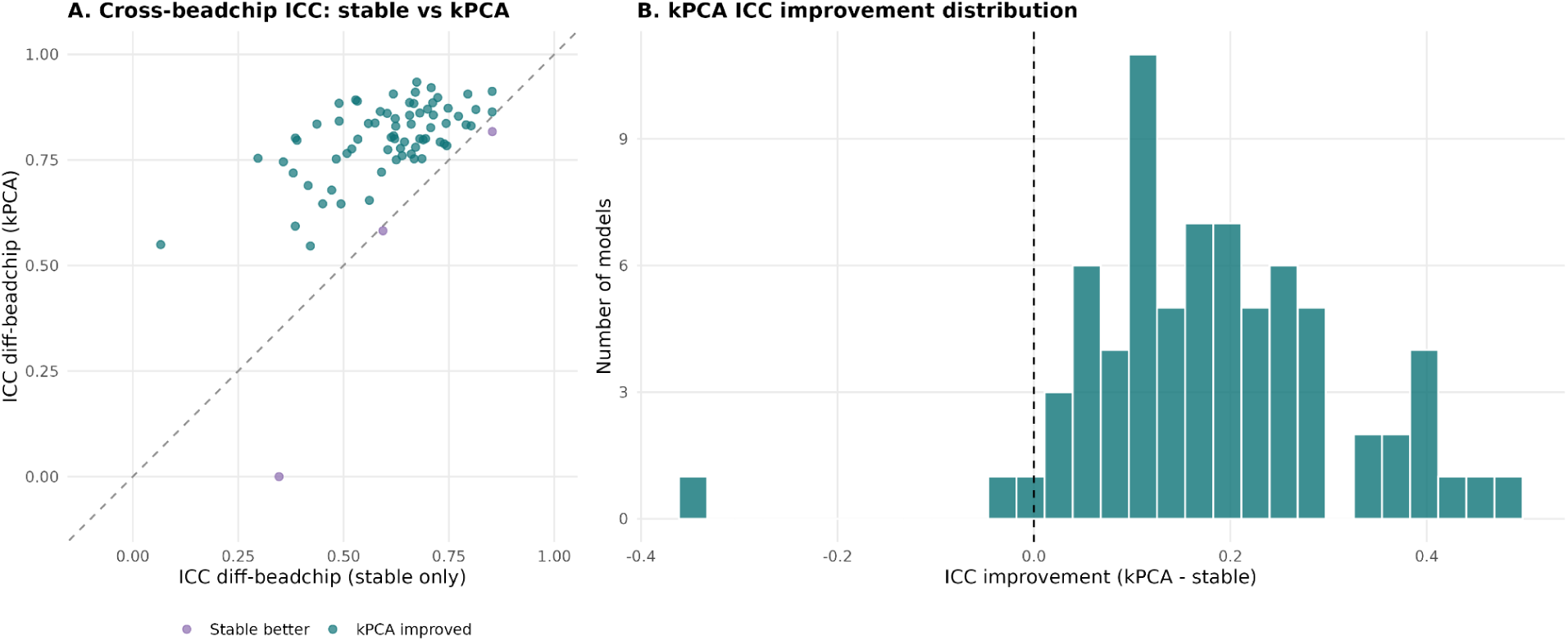
Kernel principal component analysis (kPCA) improves different-beadchip reproducibility. **(A)** Scatter plot comparing cross-beadchip ICC(2,1) for stable-probe-only models (x-axis) versus kPCA models (y-axis). kPCA was selected as the final approach for 84 biomarkers, of which 72 had a comparable stable-probe baseline and are shown here. Points above the identity line (dashed) indicate improvement with kPCA (teal); points below indicate the stable-probe version performed better (purple). **(B)** Histogram of ICC(2,1) improvement (kPCA minus stable-probe-only). Of the 72 models with a comparable stable-probe baseline, 70 (97%) showed improved cross-beadchip ICC(2,1) with kPCA, with a median improvement of +0.18 (dashed line at zero).

By integrating stable-probe feature selection with kPCA-based sample-level harmonization, we developed a robust modelling framework that maximises cross-array reproducibility. EBPs were selected using a composite score (Score = rho + 0.1 x ICC_same + 0.5 x ICC_diff; Methods), benchmarked against target criteria of rho > 0.35, same-beadchip ICC(2,1) > 0.70, and different-beadchip ICC(2,1) > 0.51; 134 models were retained. Nicotinamide and docosapentaenoic acid were excluded because they did not meet the criteria. Nicotinamide had different-beadchip ICC(2,1) = 0, which further inspection attributed to prediction-variance collapse in out-of-training samples and docosapentaenoic acid failed both the stable-probe and kPCA validity criteria, with a lower composite score than retained panels. All 134 EBPs were trained natively on MSA data: for 132, the best-performing iteration across feature-selection thresholds and kPCA configurations was selected, whereas Cysteine and Leptin performed best in an earlier MSA-trained iteration that was retained. With ICC(2,1) and rho values rounded to two decimal places, all 134 models (100%) achieved same-beadchip ICC(2,1) >= 0.80 (median 0.97; range 0.80-1.00), 129 (96.3%) achieved different-beadchip ICC(2,1) >= 0.60 (median 0.81; range 0.54-0.99), 117 (87.3%) showed Spearman rho >= 0.40, and 133 (99.3%) showed rho >= 0.35 (median 0.48; range 0.34-0.93) (Figure 4). The final models comprised 79 ElasticNet, 53 LightGBM, and 2 RandomForest models; 84 selected the kPCA pathway, 48 the stable-probe-only pathway, and 2 (Cysteine, Leptin) retained an earlier MSA-trained iteration, reflecting the fact that the impact of nonlinear batch structure varied across endpoints.

**Figure 4.**
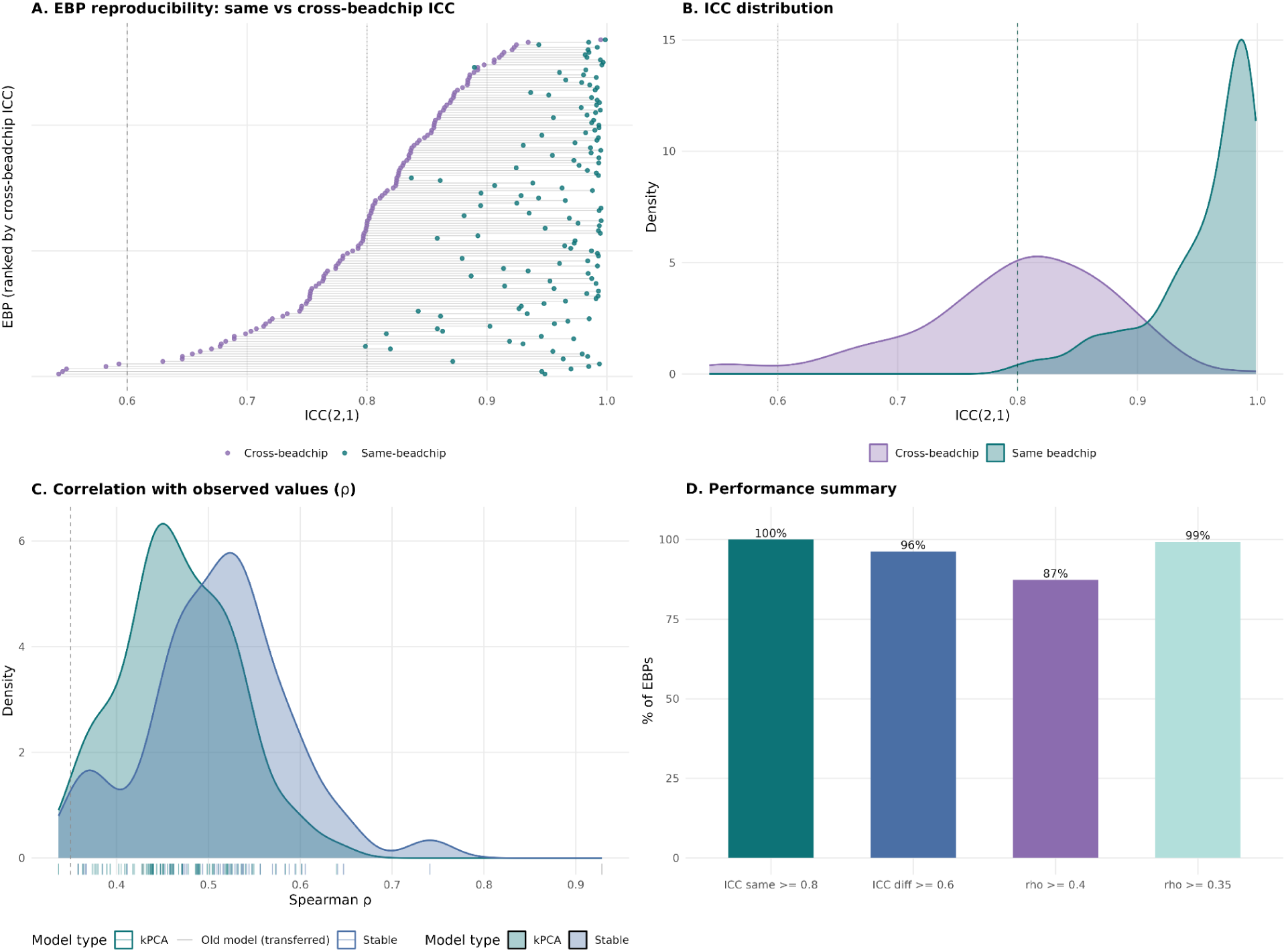
Performance of 134 MSA epigenetic biomarker proxies (EBPs). **(A)** Horizontal dot plot showing same-beadchip (teal) and cross-beadchip (purple) ICC(2,1) for each EBP, ranked by cross-beadchip ICC. Grey connecting segments illustrate the gap between same- and different-beadchip reproducibility for each model. Dashed line = ICC 0.6; dotted line = ICC 0.8. **(B)** Density plot of ICC(2,1) distributions for same-beadchip (teal) and different-beadchip (purple) comparisons, showing the separation between same-chip and different-beadchip reproducibility. **(C)** Density plot of Spearman rho (correlation between MSA-estimated and observed values), stratified by selected model type: kPCA (teal) versus stable-probe only (slate blue). Rug marks along the x-axis indicate individual EBPs. **(D)** Summary bar plot showing the percentage of EBPs meeting key performance benchmarks: same-beadchip ICC(2,1) ≥ 0.8 (100%), different-beadchip ICC(2,1) ≥ 0.6 (96.3%), Spearman rho ≥ 0.4 (87.3%), and rho ≥ 0.35 (99.3%). Final models comprised 79 ElasticNet, 53 LightGBM, and 2 RandomForest, with 84 selecting the kPCA pathway, 48 the stable-probe-only pathway, and 2 (Cysteine, Leptin) retaining an earlier MSA-trained iteration.

### Transferring Epigenetic Clocks to the MSA Platform

Building on the stable-probe and kPCA harmonization strategies, we next evaluated how existing epigenetic clocks could be transferred to the MSA platform. Because each clock differs in its modelling structure, training target, and probe composition, we adopted clock-specific approaches to ensure compatibility across Illumina arrays while maintaining the expected measurement scale.

#### Directly retrained clocks: OMICmAge and SystemsAge

OMICmAge and SystemsAge were implemented using a train-to-estimate approach, whereby the models were retrained to reproduce the biological age estimates generated by the original models rather than directly transferring their probe weights. Using paired EPICv1–MSA samples, we first applied the original models to the EPICv1 data to obtain reference biological age estimates. These EPICv1-derived estimates were then used as the target outcomes to train new models using the corresponding MSA methylation data. Model training was restricted to probes demonstrating high technical reproducibility on the MSA platform, defined as ICC ≥ 0.75 and MAE < 0.05 across same-beadchip MSA replicates.

OMICmAge achieved strong agreement between EPICv1-observed and MSA-estimated values in held-out test samples (rho = 0.944, ICC(2,1) = 0.948, n = 432; Figure 5), with high technical reproducibility across replicates (same-beadchip ICC(2,1) = 0.993, different-beadchip ICC(2,1) = 0.983). SystemsAge total and organ-specific sub-clocks showed comparable transfer accuracy in the same test samples (rho = 0.912–0.949, ICC(2,1) = 0.886–0.952; Figure 5) and replicate stability (same-beadchip ICC(2,1) = 0.976–0.995, different-beadchip ICC(2,1) = 0.935–0.975). These results demonstrate that MSA provides a suitable platform for age-related epigenetic estimators, with negligible loss of precision relative to the source arrays.

**Figure 5.**
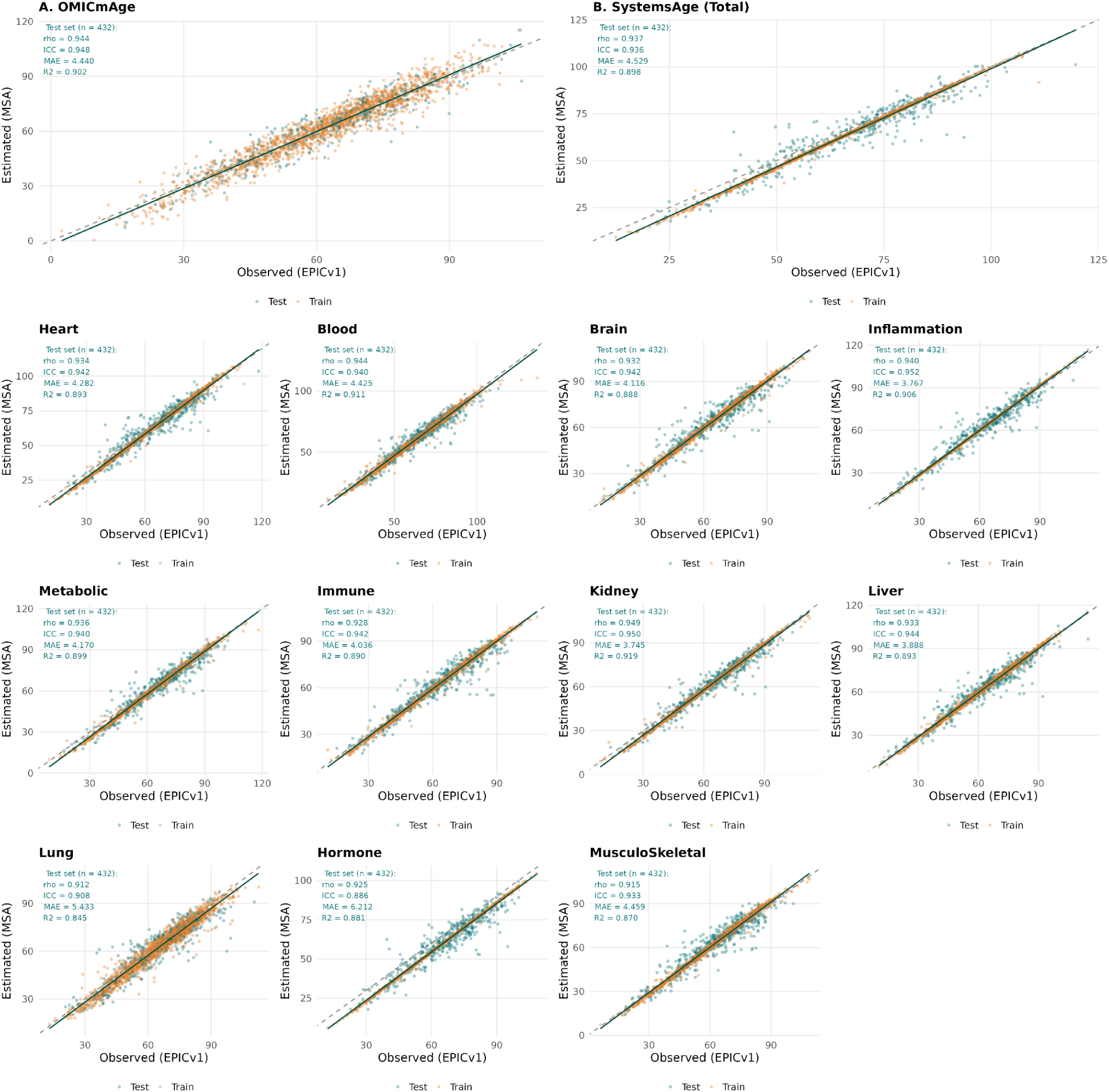
Epigenetic clock transfer to MSA: OMICmAge and SystemsAge. Scatter plots of observed (EPICv1-derived) versus MSA-estimated values for OMICmAge **(A)** and SystemsAge total **(B)**, followed by 11 organ-specific SystemsAge sub-clocks: Heart, Blood, Brain, Inflammation, Metabolic, Immune, Kidney, Liver, Lung, Hormone, and MusculoSkeletal. Points are coloured by train (amber) and test (teal; n = 432) splits. Each panel is annotated with Spearman rho, ICC(2,1), MAE, and R-squared computed on the held-out test set. Dashed line = identity; solid line = linear fit. OMICmAge was trained using ElasticNet; all SystemsAge models used LightGBM.

#### Normalization-based transfer: DunedinPACE

In contrast to OMICmAge and SystemsAge, DunedinPACE required a calibration step because the clock depends on highly consistent probe-specific scaling across platforms. Of the 173 CpGs used in the original algorithm, 171 were present on MSA. Without correction, platform-related distributional differences introduced systematic bias into the score.

To harmonize DunedinPACE across arrays, we employed a two-step strategy. First, we utilized the 40,786 probes with high reproducibility between EPICv1 and MSA (MAE < 0.05 and ICC(2,k) > 0.6). These stable probes served as anchors for aligning methylation distributions across platforms, correcting global scaling differences while preserving individual probe relationships. Second, a robust linear calibration model was fitted on a training set to align MSA-derived DunedinPACE values onto the EPICv1 scale, removing residual platform bias, while preserving inter-individual rank structure.

Performance was assessed by splitting 1,764 EPICv1 - MSA paired samples into three equal sets (n = 588 each): one for calibration training (Set 1) and two independent test sets (Sets 2 and 3). The MSA-native estimate (MSAPACEa) showed strong rank-order agreement with EPICv1 DunedinPACE (Spearman rho = 0.886–0.928 across sets) but exhibited systematic offset (MAE = 0.085–0.089; ICC(2,1) = 0.784–0.810). After robust linear calibration, cross-platform agreement improved substantially, whereby ICC increased to 0.891-0.925, MAE decreased to 0.041-0.050, and Spearman rho was preserved at 0.886-0.928 (R^2^ = 0.795-0.856) across all three sets, including both independent test sets (Figure 6). This combined workflow allowed DunedinPACE to achieve cross-array agreement comparable to the EPICv1 platform.

**Figure 6.**
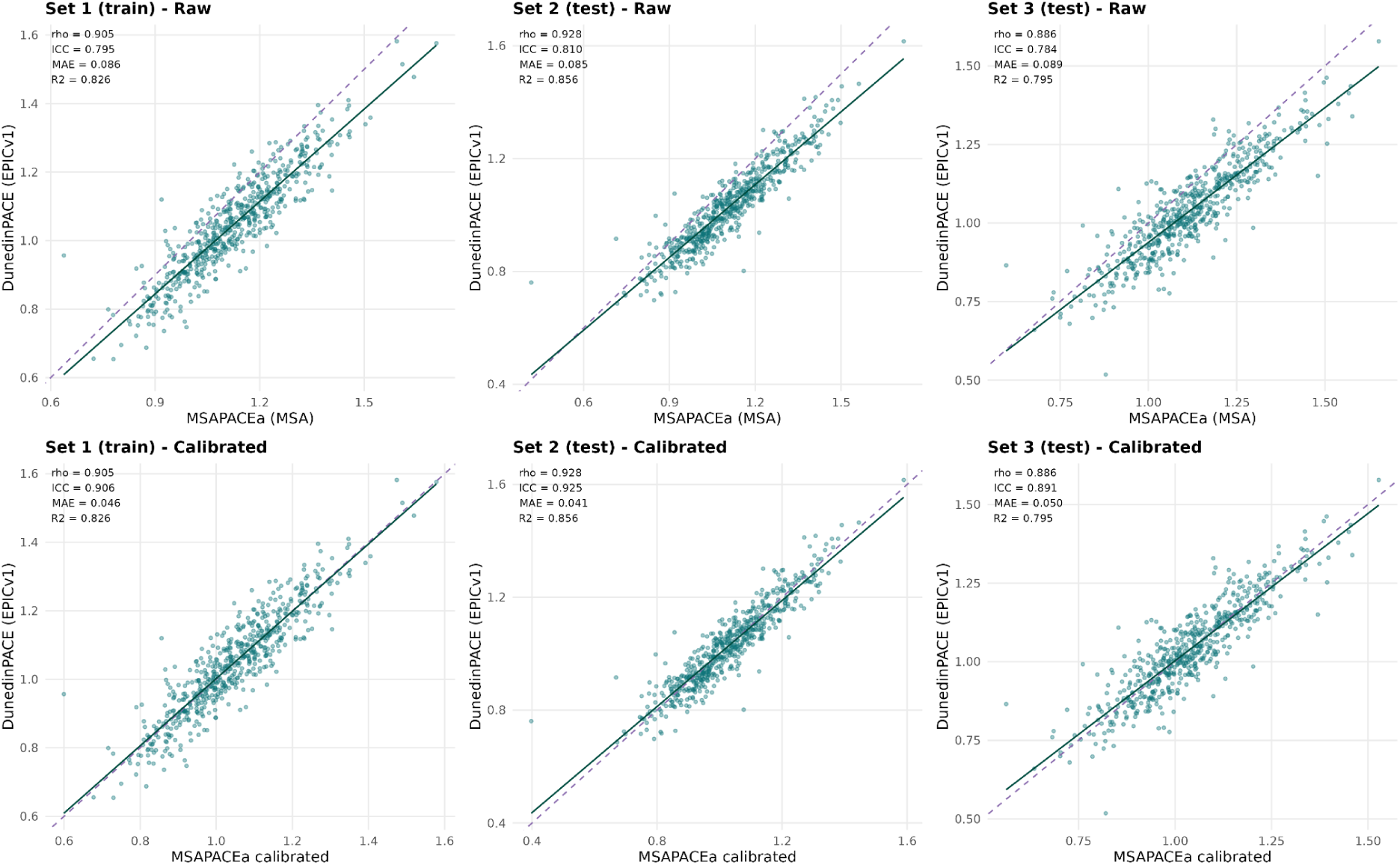
DunedinPACE transfer to MSA: normalisation and linear calibration (N = 1,764 paired samples). Scatter plots comparing MSA-derived DunedinPACE (MSAPACEa) with EPICv1-derived DunedinPACE across three independent sample sets (n = 588 each; random split). **Top row:** Pre-calibration (raw MSAPACEa) showing strong rank-order agreement (Spearman rho = 0.905, 0.928, 0.886 for Sets 1-3) but systematic offset from the identity line (dashed purple; MAE = 0.085-0.089; ICC(2,1) = 0.784-0.810). **Bottom row:** Post-calibration after robust linear regression (trained on Set 1 and applied to Sets 2 and 3), showing improved absolute agreement (ICC(2,1) = 0.906, 0.925, 0.891; MAE = 0.046, 0.041, 0.050) with preserved rank-order structure. Each panel is annotated with Spearman rho, ICC(2,1), MAE, and R-squared. Teal points; solid line = linear fit; dashed purple line = identity.

## Methods

### Study cohort and sample design

Biological replicates were obtained from the Harvard Massachusetts General Brigham (MGB) Biobank and TruDiagnostic clinical cohorts. DNA methylation was profiled on two Illumina array platforms: the Infinium MethylationEPIC BeadChip (EPICv1; ∼866,000 CpG sites) and the Infinium Methylation Screening Array (MSA; ∼284,000 CpG sites). Paired samples across platforms enabled direct cross-array comparison of probe-level methylation values. The study included within-array replicates (same and different beadchips for MSA) and cross-array replicates (EPICv1-MSA), as detailed in Supplementary Table 1. The primary cross-array comparison comprised 1,764 EPICv1-MSA paired samples from the Harvard MGB cohort, after removal of QC-failing samples.

### DNA methylation preprocessing

Raw IDAT files were preprocessed using the SeSAMe Bioconductor package (v1.18+) with the single-sample noob (ssNoob) normalisation pipeline, which performs background subtraction and dye-bias correction on a per-sample basis. For all arrays, we calculated p-value detection using pOOBAH and masked probes with p-value > 0.05. For MSA, replicate probes were collapsed to 10-character CpG prefixes using SeSAMe’s betasCollapseToPfx() function. For within-array MSA replicate analyses, uncollapsed probe identifiers were retained.

### Probe stability assessment

Cross-array probe reproducibility was quantified using two complementary metrics computed on paired biological replicates (Supplementary Table 2). Mean absolute error (MAE) was calculated as the average absolute difference in beta values between matched samples across platforms, providing a scale-dependent measure of measurement disagreement. Intraclass correlation coefficients were computed using two-way random-effects models with absolute agreement: ICC(2,1) for single measurements and ICC(2,k) for the mean of k measurements, implemented via the *irr* R package. ICC(2,1) is appropriate for generalising single-measurement comparisons across platforms, while ICC(2,k) reflects expected stability when averaged measurements are used.

Probes were classified into three stability tiers based on joint thresholds: stable (MAE < 0.05 and ICC(2,k) > 0.6), intermediate (one criterion met), and unstable (neither criterion met). For visualisation, probes were additionally classified into four ICC tiers: Poor (ICC < 0.4), Fair (0.4 <= ICC < 0.6), Good (0.6 <= ICC < 0.75), and Excellent (ICC >= 0.75). Stability metrics were computed for three array comparisons (Table 1): EPICv1 - MSA and MSA within-array replicates on the same and on different beadchips.

#### Epigenetic biomarker panel (EBP) development

##### Stable-probe feature selection

For EBP model training, the feature space was restricted to probes demonstrating high same-beadchip reproducibility in MSA same-beadchip technical replicates (ICC(2,1) ≥ 0.75 and 0.05 < mean beta < 0.95). This filtering step retained 99,378 MSA probes with minimal technical noise for downstream modelling.

##### Kernel principal component analysis (kPCA) harmonization

To address residual nonlinear batch effects across beadchips that persisted after probe-level filtering, kernel principal component analysis (kPCA) was applied to the stable-probe methylation matrix prior to model training. kPCA maps samples into a higher-dimensional feature space via a kernel function. By merging signal across correlated probes, reducing collinearity, and lowering effective dimensionality, this transformation can reduce sensitivity to probe-specific technical noise, though separation of platform-driven variation from biological signal is not guaranteed a priori and was evaluated empirically for each biomarker. For each biomarker, models were trained on both the stable-probe features directly and on kPCA-transformed features and the one with the highest performance was selected

##### Model training and selection

EBP models were trained using three machine learning frameworks: ElasticNet (penalised linear regression), LightGBM (gradient-boosted decision trees), and RandomForest. Target values for each biomarker were derived from observed clinical, proteomic, or metabolomic measurements. Model performance was evaluated using Spearman rank correlation (rho) between predicted and observed values, and reproducibility was assessed using ICC(2,1) computed on same-beadchip and different-beadchip MSA technical replicates.

For each biomarker, a composite score, Score = rho + 0.1 x ICC_same + 0.5 x ICC_diff - where rho is the Spearman correlation between MSA-estimated and observed values and ICC_same and ICC_diff are the same- and different-beadchip ICC(2,1) - was computed for both the stable-probe and kPCA versions, and the higher-scoring version was selected as the final model. The composite score was the primary selection criterion; candidate panels were additionally benchmarked against target criteria of rho > 0.35, ICC_same > 0.70, and ICC_diff > 0.51, which the retained panels generally satisfied (Results). All biomarkers were trained on MSA data; for Cysteine and Leptin, an earlier MSA-trained iteration outperformed subsequent iterations and was retained as the final model. Nicotinamide was excluded due to different-beadchip ICC(2,1) = 0, which inspection attributed to prediction-variance collapse on out-of-training samples (53% missing observed values in training). Docosapentaenoic acid was excluded for failing both the stable-probe and kPCA validity criteria with a lower composite score than the retained panels. The final panel comprises 134 EBPs.

#### Epigenetic clock transfer

##### OMICmAge and SystemsAge

OMICmAge and SystemsAge (comprising a total score and 11 organ-specific sub-clocks) are train-to-estimate clocks: rather than transferring probe weights directly, the models were retrained on the MSA platform to predict their target epigenetic age estimates derived from the source array (EPICv1). Training used probes identified as stable between MSA same-beadchip replicates (Supplementary Table 1) as input features and the EPICv1-derived clock values as targets. The best model for all SystemsAge metrics was based on LightGBM, while it was ElasticNet for OMICmAge.

After model training, two sequential linear corrections were applied to align MSA-derived estimates with their EPICv1 reference values. For OMICmAge, a post-training MSA-to-EPIC correction was computed on held-out test samples by fitting the slope and intercept of the MSA estimate against the EPICv1 estimate, and applying the inverse linear transform: corrected = (raw - intercept) / slope. For all SystemsAge metrics, an age correction was applied using slope and intercept values fitted against chronological age in a large balanced reference cohort (Harvard + TruD), using the same inverse linear formula. These corrections remove residual platform-related scaling differences while preserving inter-individual rank structure.

Model performance was assessed using Spearman rho between MSA-estimated and EPICv1-observed values, and reproducibility was quantified using ICC(2,1) on same-beadchip and different-beadchip MSA replicates.

##### DunedinPACE

DunedinPACE was transferred to the MSA platform using a two-step normalisation and calibration procedure rather than retraining, because the clock relies on specific probe-weight scaling that must be preserved across platforms.

### Step 1: Stable-probe normalisation

Of the 173 CpGs used in the original DunedinPACE algorithm, 171 were present on the MSA array. To correct platform-related distributional differences, we identified 40,786 CpGs with high cross-array reproducibility between EPICv1 and MSA (MAE < 0.05 and ICC(2,k) > 0.6). These stable probes served as normalisation anchors: their distributions were used to align the MSA methylation scale to the EPICv1 reference, correcting global scaling differences while preserving probe-level relationships. This normalisation step was implemented in the ‘MSAPACÈ function (MSAPACEa variant, using 40,786 normalisation probes). The normalised MSA beta values were then passed through the original DunedinPACE ‘PACEProjector’ algorithm to generate MSA-derived DunedinPACE scores. Unlike the same-beadchip MSA replicate filtering used for the EBP and clock feature selection, DunedinPACE normalisation required probes stable across platforms, since the anchors must reflect the specific comparison being corrected, cross-array agreement. As such, we used the ICC(2,k) > 0.6 threshold to reflect the variance of the cross-array comparisons.

### Step 2: Linear calibration

After normalisation, residual platform bias in the DunedinPACE score was corrected using robust linear regression using the *rlm* function from the MASS package. The 1,764 EPICv1-MSA paired samples were randomly split into three equal sets (n = 588 each). Set 1 served as the calibration training set: a robust linear model was fitted with MSAPACEa as the predictor and EPICv1-derived DunedinPACE (computed using SeSAMe with manual pOOBAH masking) as the response. The fitted intercept (a) and slope (b) were then applied as a forward prediction (corrected = a + b * MSAPACEa) to Sets 2 and 3 as independent validation. Note that this calibration direction differs from the inverse correction used for the train-to-estimate clocks above, because the rlm model was fitted to predict the EPICv1 reference from the MSA value directly.

#### Statistical analysis

Spearman rank correlation coefficients were used to assess monotonic agreement between predicted and observed values. Intraclass correlation coefficients (ICC(2,1), two-way random-effects, absolute agreement) were computed using the *irr* R package to quantify measurement reproducibility. MAE was computed as the mean of absolute pairwise differences. R-squared values were obtained from ordinary least squares linear regression. For the kPCA improvement analysis, the change in different-beadchip ICC (kPCA minus stable-only) was computed for each model where kPCA was selected. Analyses were performed in R (v4.3+) using the Bioconductor *SeSAMe*, *irr*, *MASS* packages, as well as in Python using Python 3.11. Machine-learning models were implemented using scikit-learn v1.6.1 and LightGBM v4.5.0, with hyperparameter optimization conducted using Optuna v4.3.0. Model interpretation was performed using SHAP v0.45.0 and FastTreeSHAP v0.1.6. Numerical computation and data processing were supported by NumPy v1.26.4, SciPy v1.17.1, pandas v2.2.3, and Polars v1.22.0.

## Discussion

In this study, we developed and validated a generalizable, reproducibility-centered framework for transitioning DNAm algorithms from legacy Illumina EPIC-based platforms to the Illumina Methylation Screening Array (MSA). Leveraging paired EPICv1 - MSA and MSA - MSA samples, we systematically identified reproducible CpG sites using complementary measures of technical agreement and used this stable probe set to support two analytical streams: the development of 134 MSA-compatible epigenetic biomarker proxies (EBPs) and the transfer of established epigenetic clocks. For biomarker development, we compared models trained on stable probes alone to those incorporating kernel principal component analysis (kPCA) for nonlinear harmonization, selecting final models based on performance and replicate reproducibility. For clock transfer, we applied model-specific strategies, including retraining of OMICmAge and SystemsAge ^5,15^ and normalization plus calibration for DunedinPACE ^3^. Across all applications, we demonstrate strong cross-platform agreement and high same- and different-beadchip reproducibility, establishing a unified framework for deploying DNAm algorithms on the MSA platform.

A central finding of this study is that cross-array probe overlap alone is insufficient to ensure reliable biomarker performance, highlighting the importance of explicitly quantifying probe-level reproducibility. While prior work has often relied on intersecting CpG sets, direct transfer of legacy models, or limited cross-platform validation approaches ^9,10,16^, our results demonstrate that a substantial fraction of shared probes exhibit poor cross-platform agreement despite being present on both arrays. By jointly considering absolute agreement and rank-order consistency, we show that probe quality can be systematically stratified, enabling the definition of a stable feature space that preserves biological signal while minimizing technical noise. This provides a principled alternative to ad hoc probe filtering approaches and suggests that reproducibility-based feature selection should become a standard prerequisite for cross-platform DNAm analyses. More broadly, our findings reinforce the notion that CpG presence alone is not sufficient for biomarker portability; rather, reproducible measurement characteristics are essential for preserving model performance across platforms.

Our results further demonstrate that probe-level filtering, while necessary, is not sufficient to fully address technical variation. Even after restricting models to stable CpGs, residual nonlinear effects related to beadchip and platform structure persisted and degraded reproducibility for a subset of biomarkers. The application of kPCA provided a consistent and substantial improvement in cross-beadchip agreement, particularly for models with lower baseline stability, without compromising biological validity. These findings extend prior normalization and batch-correction approaches, including ComBat, functional normalization, and related harmonization frameworks ^17–19^, by demonstrating that nonlinear sample-level harmonization can complement reproducibility-informed feature selection to better isolate biological signal from technical variation. Importantly, the benefits of kPCA were most pronounced among biomarkers that initially exhibited poorer reproducibility, suggesting that residual nonlinear structure may represent an underappreciated source of variability in DNAm biomarker development. Collectively, these results highlight the importance of addressing both feature-level and sample-level sources of variation when deploying DNAm biomarkers across platforms.

Another key insight is that different classes of DNAm algorithms require distinct strategies for cross-platform transfer. Consistent with their design, train-to-estimate clocks such as OMICmAge and SystemsAge ^5,15^ could be effectively retrained on MSA to reproduce EPIC-derived reference values with minimal loss of precision. In contrast, DunedinPACE, which depends on precise probe-specific scaling and calibration to preserve its intended measurement scale ^3^, could not be directly retrained and instead required normalization and calibration procedures to achieve cross-platform agreement. This distinction is often underappreciated in the literature, where cross-platform transfer is frequently treated as a uniform problem. Our results demonstrate that algorithm architecture fundamentally constrains transferability and that successful deployment requires model-specific adaptation strategies. More generally, these findings suggest that future efforts to migrate DNAm algorithms across platforms should consider the biological and statistical assumptions embedded within each model rather than relying on a single transfer methodology.

Importantly, we show that the MSA platform can support a broad range of DNAm biomarkers with high technical reproducibility and preserved biological relevance. The successful development of 134 EBPs, the majority of which achieved strong same- and different-beadchip ICC values together with meaningful correlations to observed traits, demonstrates that reduced probe content does not preclude robust biomarker performance when appropriate feature selection and harmonization strategies are applied. Similarly, the successful transfer of multiple epigenetic clocks underscores the feasibility of deploying established DNAm estimators on a lower-cost, scalable platform. These findings are particularly relevant as DNAm biomarkers continue to expand beyond observational studies into intervention trials, population-scale cohorts, and emerging clinical applications ^20,21^. Together, our results position the MSA as a practical alternative for large-scale and translational applications where cost, throughput, and reproducibility are critical considerations.

Several limitations should be considered. First, although we leveraged large paired datasets across multiple arrays, the evaluation was restricted to specific cohorts and may not fully capture variability across diverse populations, tissues, or experimental conditions. Second, while kPCA improves reproducibility in most cases, it introduces an additional layer of model complexity that may affect interpretability and generalizability in certain contexts. Third, our framework was evaluated on a defined set of biomarkers and clocks, and additional work will be needed to assess its applicability to other DNAm models, including those trained in different tissues or disease contexts. Finally, while the MSA shows strong performance, its reduced probe content inherently limits genomic coverage relative to EPIC arrays, which may constrain certain applications. Future work should focus on validating these approaches in independent and more diverse cohorts, exploring alternative harmonization strategies, and extending this framework to emerging methylation platforms and multi-omic biomarker systems ^22^. Despite these limitations, our results provide a robust foundation for reproducibility-aware deployment of DNAm biomarkers and a scalable path forward for cross-platform translation in epigenetic research and clinical applications.

## Supplementary Tables

**Supplementary table 1:** Biological replicates in the same and different Illumina arrays.

| Dataset | N. Samples | Replicates per sample | Arrays | Beadchips | Cohort |
| --- | --- | --- | --- | --- | --- |
| MSA same beadchip replicates | 24 | 2 | MSA | Same beadchips | Harvard MGB |
| MSA different beadchips | 32 | 2 | MSA | Different beadchips | Harvard MGB |
| MSA, EPICv1 | 1,764 | 2 | EPICv1, MSA | Different arrays | Harvard MGB |

**Supplementary Table 2:**
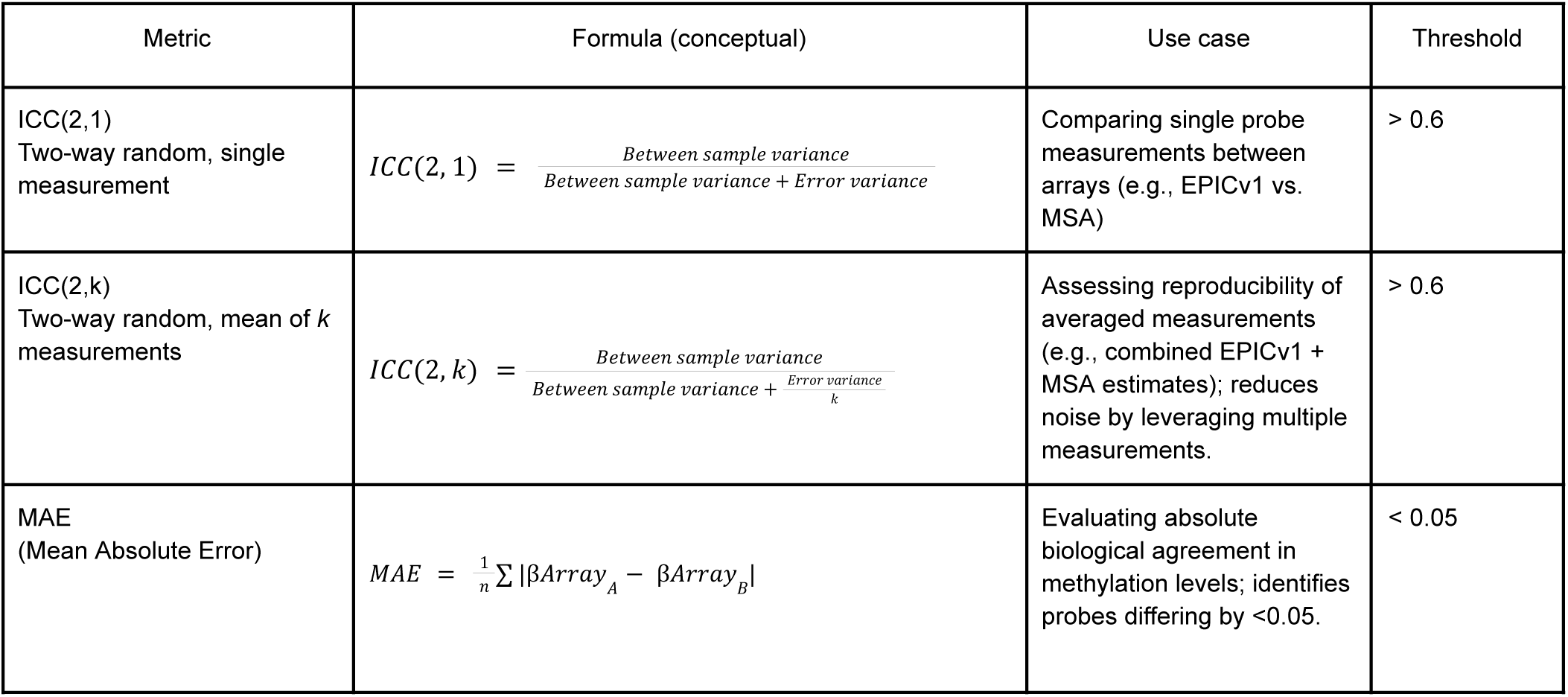
Reproducibility metrics, Formulas, Use cases, and Thresholds.

## Supplementary Figures

**Supplementary Figure 1.**
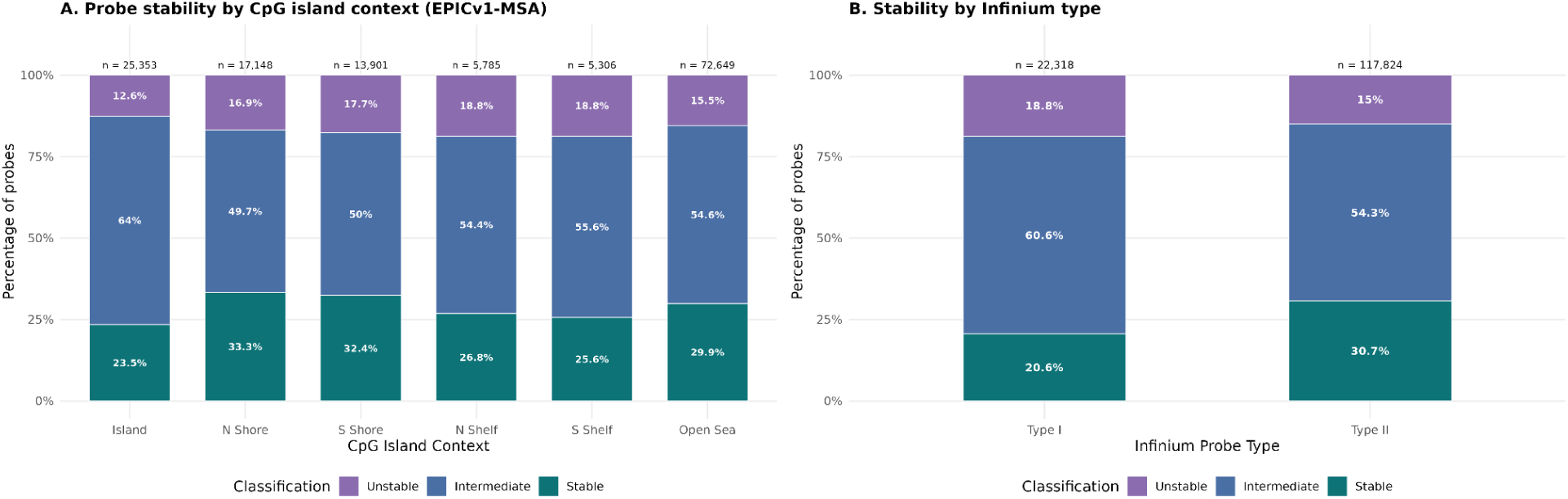
Probe stability by genomic context and Infinium type (EPICv1-MSA). **(A)** Stacked bar plot showing the proportion of probes classified as stable (MAE < 0.05 and ICC(2,k) > 0.6; teal), intermediate (one criterion met; slate blue), and unstable (neither criterion met; purple) across six CpG island contexts. Stability rates are relatively uniform: Island (23.5%), N Shore (33.3%), S Shore (32.4%), N Shelf (26.8%), S Shelf (25.6%), and Open Sea (29.9%), indicating no strong enrichment or depletion of stable probes by genomic context. Total probe counts are shown above each bar. **(B)** Same three-tier classification by Infinium probe type. Type II probes show modestly higher stability (30.7%) than Type I probes (20.6%), consistent with the simpler single-channel chemistry of Type II designs. N = 140,142 EPICv1-MSA probes with annotation.

**Supplementary Figure 2.**
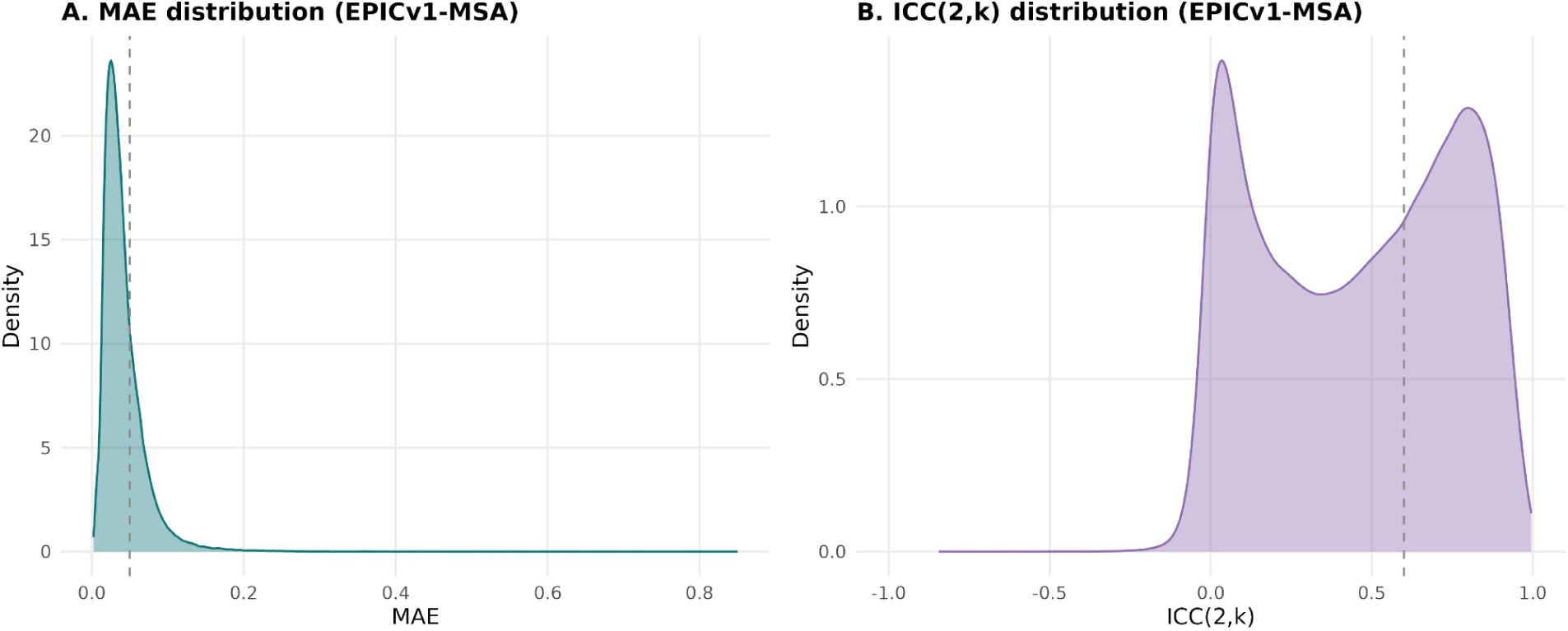
MAE and ICC distributions for EPICv1-MSA probes. **(A)** Density plot of MAE values for 140,150 shared CpG sites between EPICv1 and MSA, showing a strongly right-skewed distribution with the majority of probes having low MAE. Dashed line indicates the MAE < 0.05 stability threshold. **(B)** Density plot of ICC(2,k) values for the same probes, showing a bimodal distribution with peaks near zero (low-variance probes with poor reproducibility) and near 1.0 (highly reproducible probes). Dashed line indicates the ICC > 0.6 stability threshold.

## Notes

### Competing Interest Statement

KBS, SF, IG, LBD, VBD, MM, RS and NCG are employees of TruDiagnostic.

